# Red List criteria underestimate climate-related extinction risk of range-shifting species

**DOI:** 10.1101/2025.08.20.671260

**Authors:** Raya Keuth, Susanne A. Fritz, Damaris Zurell

## Abstract

Climate change causes global species redistribution and elevates extinction risk, making early identification of vulnerable species critical for timely conservation. The IUCN Red List provides guidelines for assessing climate-related extinction risk using species distribution models (SDMs) and spatially explicit population models (SEPMs). However, a systematic evaluation of these guidelines is currently missing. Using simulations of virtual species with diverse life-history traits and range dynamics, we found that SDMs consistently underestimated extinction risk for range-shifting species. This was due to a concave relationship between population size and habitat loss, which contradicts the linear assumption in the Red List guidelines. For range-contracting species, SDMs provided adequate warning times. Probabilistic extinction estimates from SEPMs provided delayed warning for all species, particularly for highly threatened ones. Our results reveal key limitations of current Red List guidelines under climate change. Based on our findings, we provide tentative recommendations for updating the IUCN Red List guidelines.

## Introduction

Climate change poses a major threat to biodiversity, affecting virtually all biological processes^1^ and potentially elevating extinction risk for many species^2,3^. Species dynamically respond to climate change, for example by tracking suitable habitat^4^, which has been projected to lead to substantial global redistribution of species^5^. To protect biodiversity before declines become irreversible, early identification of the most threatened species is essential. Conservation assessments should hence provide sufficient warning time, defined as the time between identification of a threatened species and its extinction without any interventions^6^.

The IUCN Red List for Threatened Species (Red List hereafter) is the most widely used framework for assessing extinction risk and informing conservation and policy decisions^7,8^. It classifies species into one of three threatened categories (Vulnerable, Endangered, Critically endangered) based on five quantitative criteria (A-E) that are derived from the theory of extinction trajectories for single populations^7,9,10^. The five different criteria provide alternative ways of assessing a species in the Red List, with qualifying for a threatened category under one criterion being sufficient to classify species in the category. The assessment of the species is mostly based on population size and trends, as well as geographic range area^9^. Originally designed to identify species already in decline, the IUCN has since introduced guidelines for assessing future extinction risk under climate change. These recommend species distribution models (SDMs) to infer habitat loss, or spatially explicit population models (SEPMs) to infer population loss or estimate extinction probability^11,12^.

Despite their growing use, the predictive performance of SDMs under climate change remains debated^13–15^. SDMs are correlative models that relate observed species occurrences to prevailing environmental conditions and use this species-environment relationship to predict the suitable habitat under future climatic scenarios^16^. However, SDMs omit key biological processes, such as demography and dispersal^17,18^, and simplistic dispersal assumptions (e.g. no dispersal or full dispersal) can lead to an over- or underestimation of the expected range size^14^. Nevertheless, SDMs currently constitute the most pragmatic approach for projecting potential future climate impacts. The Red List guidelines suggest to use SDM-predicted changes in habitat suitability as a proxy for population size reduction under criterion A3^8(p.102)^. This approach assumes a linear relationship between population loss and SDM-predicted habitat loss, which is acknowledged in the Red List guidelines as a source of uncertainty^8(p.99)^. Beyond simply causing uncertainty, this assumption actually lacks empirical validation^19,20^, and could be influenced by the non-uniform distribution of species in the landscape^21^, extinction debts^22^, and extinction thresholds related to minimum viable population sizes^23^.

SEPMs, in contrast, simulate demographic and dispersal processes over space and time, potentially offering more realistic predictions of species responses to climate change^24,25^. Yet, wider application of these models has been hampered by technical and data challenges^26,27^ and they have thus been rarely used for assessing climate-related future extinction risk of species^28^. A key open question is whether SEPMs provide more accurate or timely extinction risk estimates than SDMs, particularly for species with different range dynamics under climate change. So far, only a single case study has compared these modelling approaches in the context of IUCN assessments^29^ while a systematic evaluation of modelling approaches across varying life histories and range dynamics is missing. This gap prevents evidence-based improvements to current Red List guidelines.

In this study, we aim to test the Red List guidelines for assessing species threatened by climate change. Specifically, we test (i) whether the assumed linear relationship between habitat loss and population decline holds across species and range dynamics, and (ii) whether model-based assessments differ in their ability to identify extinction risk early enough for conservation action. Due to the lack of data regarding climate change-driven extinctions, we chose a simulation approach^30,31^. We generated sixteen virtual species with contrasting life-history and dispersal traits (niche position, niche breadth, growth rate and dispersal distance)^32^ and used a spatially explicit modelling framework^33^ to project population and range dynamics under climate change across three different stochastically replicated artificial landscapes obtained from a neutral landscape model (Fig. 1). We set up the simulations in a way that all species eventually went extinct under climate change due to habitat loss. Half of the species (those with central niche positions) first underwent a range shift before extinction, while the other half of the species (those with marginal, cold-adapted niches) only experienced range contractions, also leading to extinction. Finally, we compared the performance of SDMs and SEPMs for assessing extinction risk, focusing on their ability to provide timely warnings for species with varying life history traits and undergoing either range contraction or range shift.

**Figure 1:**
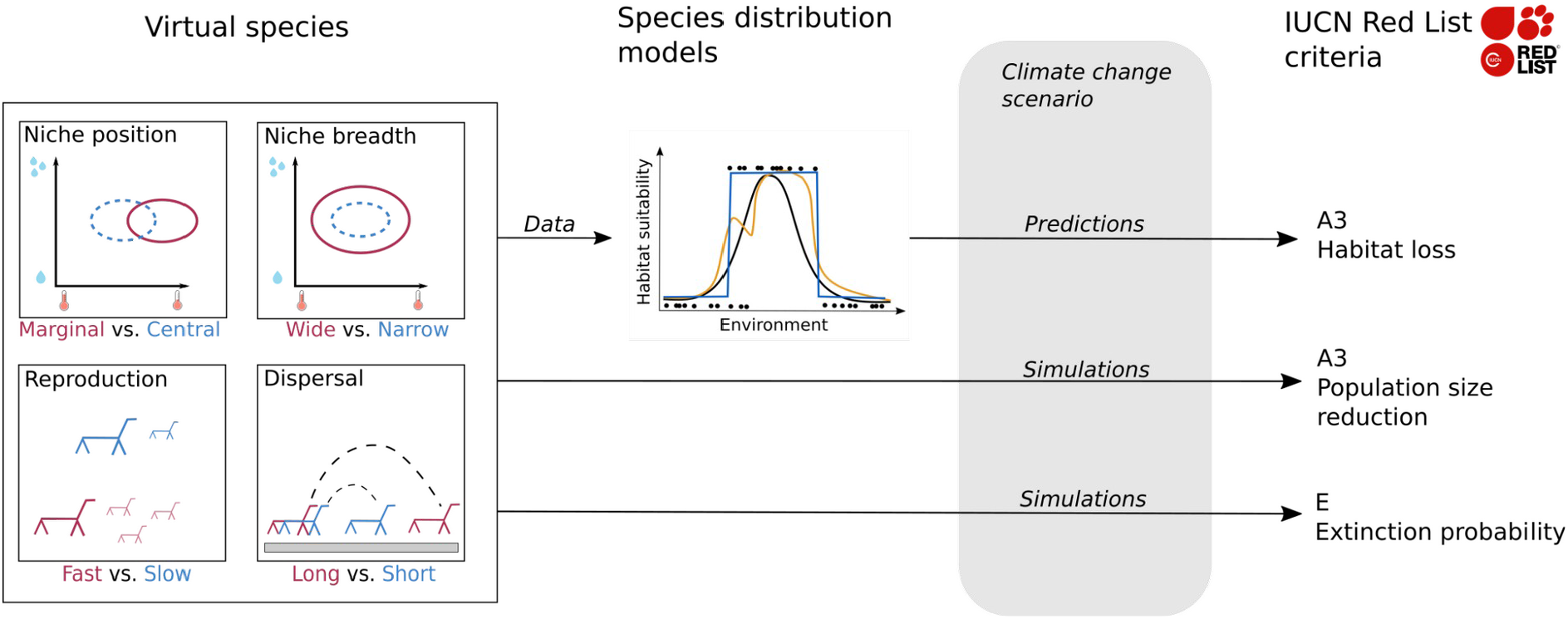
Conceptual figure of the modelling workflow. We simulated virtual species under climate change using a spatially explicit individual-based model. Species varied in four traits, which resulted in a variety of range distributions and dynamics. Before climate change set in, distribution data were sampled from the simulations as input to train correlative species distribution models (SDMs). SDMs were then used to predict habitat loss under climate change. Finally, Red List classifications under climate change were compared for SDM-derived habitat loss (criterion A3) and simulation-derived population size reduction (criterion A3) and extinction probability (criterion E).

## Results

### Population dynamics and SDM performance

The considered niche traits led to distinct differences in range and population dynamics (here, population refers to the number of individuals across the entire landscape), while the traits growth rate and dispersal only showed minor effects (Fig. S1 + S2). Notably, species with marginal niche position mainly occurred at the “northern” edge of the landscape and their ranges contracted towards that edge under climate change (Fig. 2 + Fig. S3). Species with central niche position had centrally located ranges and exhibited range shifting before going extinct under climate change (Fig. 2 + Fig. S4). Notably, not all range-shifting species (i.e., species with a central niche position) were able to fully shift their ranges and reach the northern boundary of the landscape. Some species became extinct earlier due to dispersal limitations, as suitable habitat was still available in the northern part of the landscape (Fig. 2 + Fig. S1). As could be expected, species with slow growth, narrow niches and short dispersal went extinct earlier than species with fast growth, wide niches and long dispersal (Fig. S1). The differences in temporal dynamics between species with different trait values were more pronounced for range-shifting species than for range-contracting species.

**Figure 2:**
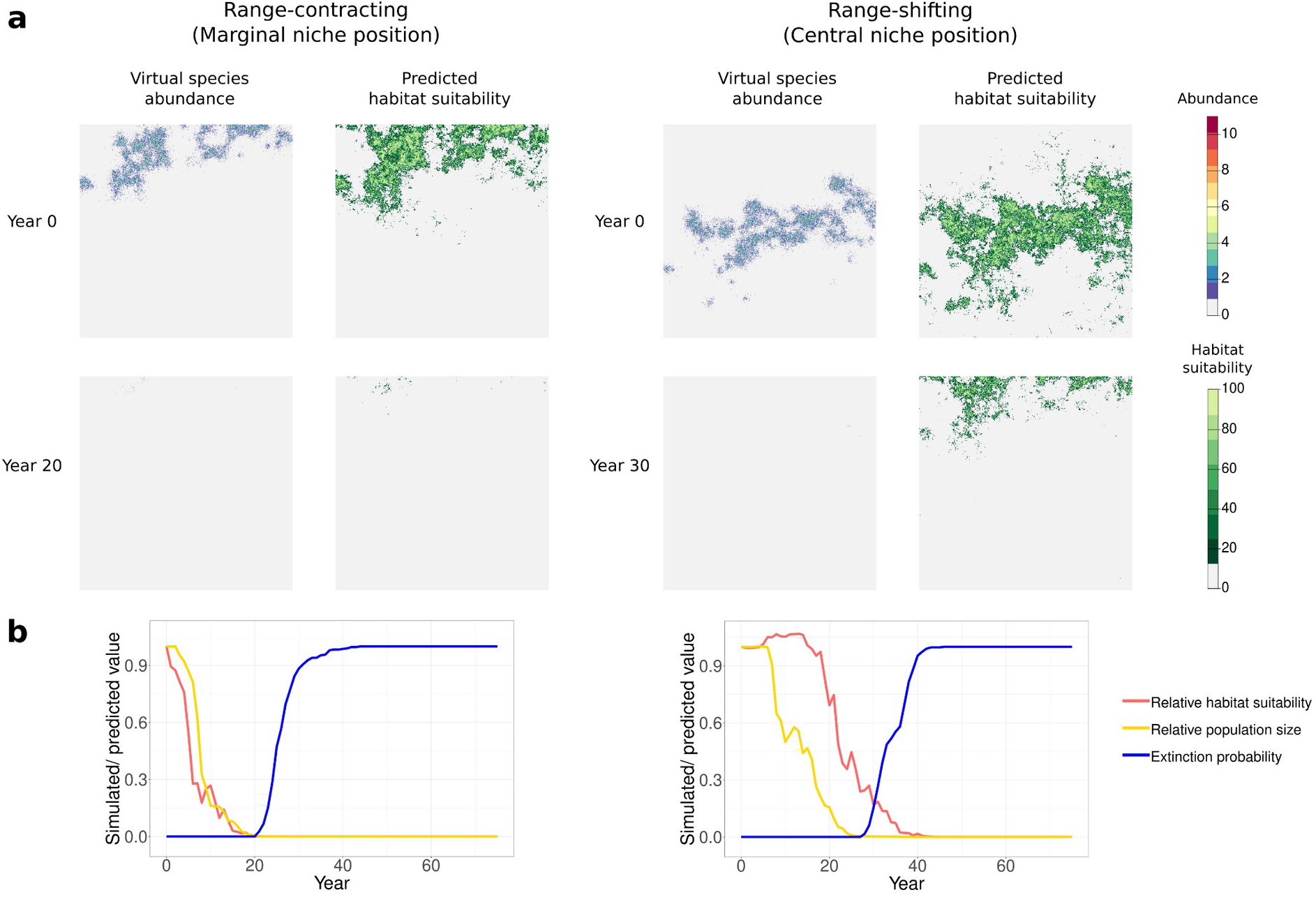
Distribution of virtual species and SDM-predicted habitat suitability for two example species. Top panels (a) show the maps of simulated „true” species abundance and maps of habitat suitability predicted by SDMs for one replicate run under equilibrium conditions (year 0) and under climate change (year 20 / 30) in one example landscape. Bottom panels (b) show the corresponding time trajectories of the relative change in habitat suitability sums predicted from SDMs and in simulated population size, both averaged over ten selected replicate runs, as well as simulated extinction probability. Results are shown for two example species differing in their range dynamics (i.e., in marginal vs. central niche position). Both species have a narrow niche, slow growth rate, and short dispersal.

We sampled species occurrence from the simulations before climate change set in, and fitted three different SDM algorithms and an ensemble model. The SDM ensembles showed high pre-climate change predictive accuracy for all species and landscapes (AUC: 0.95 ± 0.02; TSS: 0.78 ± 0.06). The predictive performance was slightly lower for species with central compared to marginal niches and for species with short compared to long dispersal (Fig. S5 + S6). Under climate change, SDMs correctly predicted range loss of range-contracting species (i.e., the distribution of simulated abundances and the increase of extinction probability aligned well with the SDM-derived habitat predictions), but they underestimated the range loss of dispersal-limited range-shifting species (i.e., the distribution of simulated abundances was smaller than the SDM-derived habitat distribution and extinction probability increased although SDMs still predicted suitable habitat; Fig. 2). This pattern was consistent across replicate landscapes (Fig. S1).

### Population size - habitat loss relationship

For all species and replicate landscapes, the relationship between relative population size and SDM-derived habitat loss observed across simulated species and landscapes deviated from the linear assumption proposed in the Red List guidelines (Fig. 3, S7 + S8). For range-contracting species (with marginal, cold-adapted niche positions), the relationship was convex with an initial smaller rate of decline of population size in comparison to habitat loss. Population size steeply declined once habitat loss exceeded 50% across all replicate landscapes (Fig. S7 + S8). In contrast, range-shifting species (with central niche positions) showed a concave relationship with considerable declines in population size already happening at low rates of habitat loss (Fig. 3). Yet, we also found considerable differences in the population size - habitat loss relationship for range-shifting species in different replicate landscapes, with some being closer to a linear relationship (Fig. S7 + S8). We only found comparatively small differences in the population size - habitat loss relationships for the other traits, with the variability between the different landscapes often being larger than among the different trait levels (Fig. S7 + S8). We quantified the relative importance of the different traits using a Bayesian generalised mixed-model analysis of covariance. This revealed that the population size - habitat loss relationship significantly differed across all traits, but the effect sizes for niche position (i.e., for range-shifting vs. range-contracting species) were considerably larger than for all other traits (Table S1).

**Figure 3:**
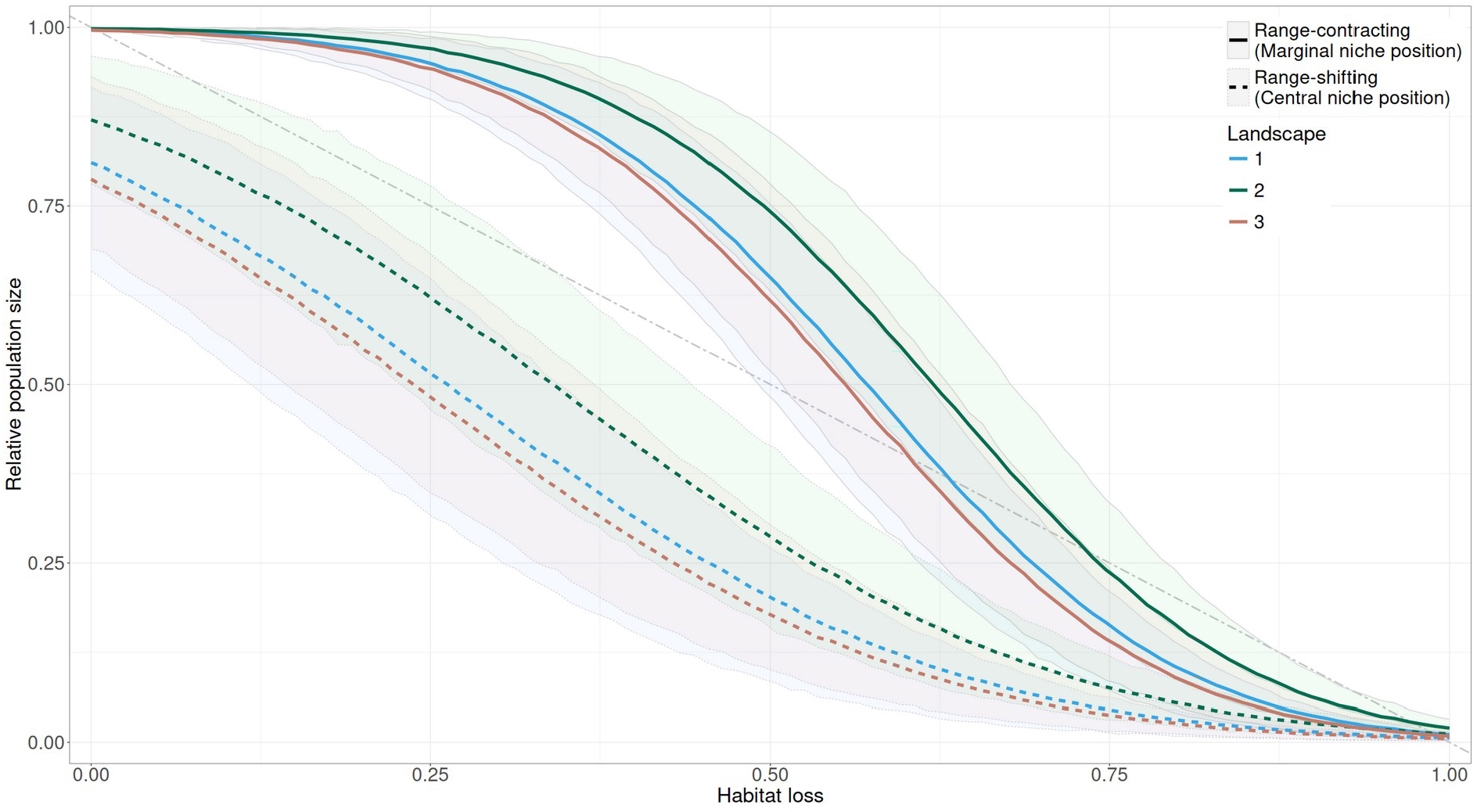
Population size - habitat loss relationship for the different range dynamics. Relative population size was obtained from the simulation (observed true data for virtual species) and habitat loss was predicted by the SDMs. The dot-dash grey line shows the assumption of a linear relationship between population size and habitat loss as proposed by the Red List guidelines. The relationships are displayed using an ordered beta regression model, which was fitted using the “true” population size values obtained from the simulations as response variable and habitat loss, the 16 trait combinations across the 10 selected replicate runs as predictor variables, and landscape as a random effect. Lines indicate the mean model predictions and the transparent bands the 95% confidence interval. We only show results here for the trait niche position, resulting in range-contracting and range-shifting species, while results for the other traits are displayed in Fig. S7.

### Red List classification

An important question is to which extent SDM-derived habitat loss under climate change can be used for assessment in the Red List. To answer this, we determined the time at which a species would first be classified into each threatened category of the Red List: VU, EN, and CR (Vulnerable, Endangered, and Critically Endangered). Specifically, we compared assessments against criterion A3 when i) applied to the true (simulated) population size and ii) to the sum of habitat suitability derived from SDMs as recommended by the Red List guidelines^8(pp.101-111)^. Additionally, we compared these to the assessment under criterion E based on the quantitative extinction probability extracted from our simulation. Since all species went extinct during the 90 years of climate change in our simulations, all of them qualified for all threatened categories throughout their life span. Again, we found the strongest differences between range-contracting and range-shifting species, reflecting the differences in the population size - habitat loss relationship (Fig. 4 + Fig. S9).

**Figure 4:**
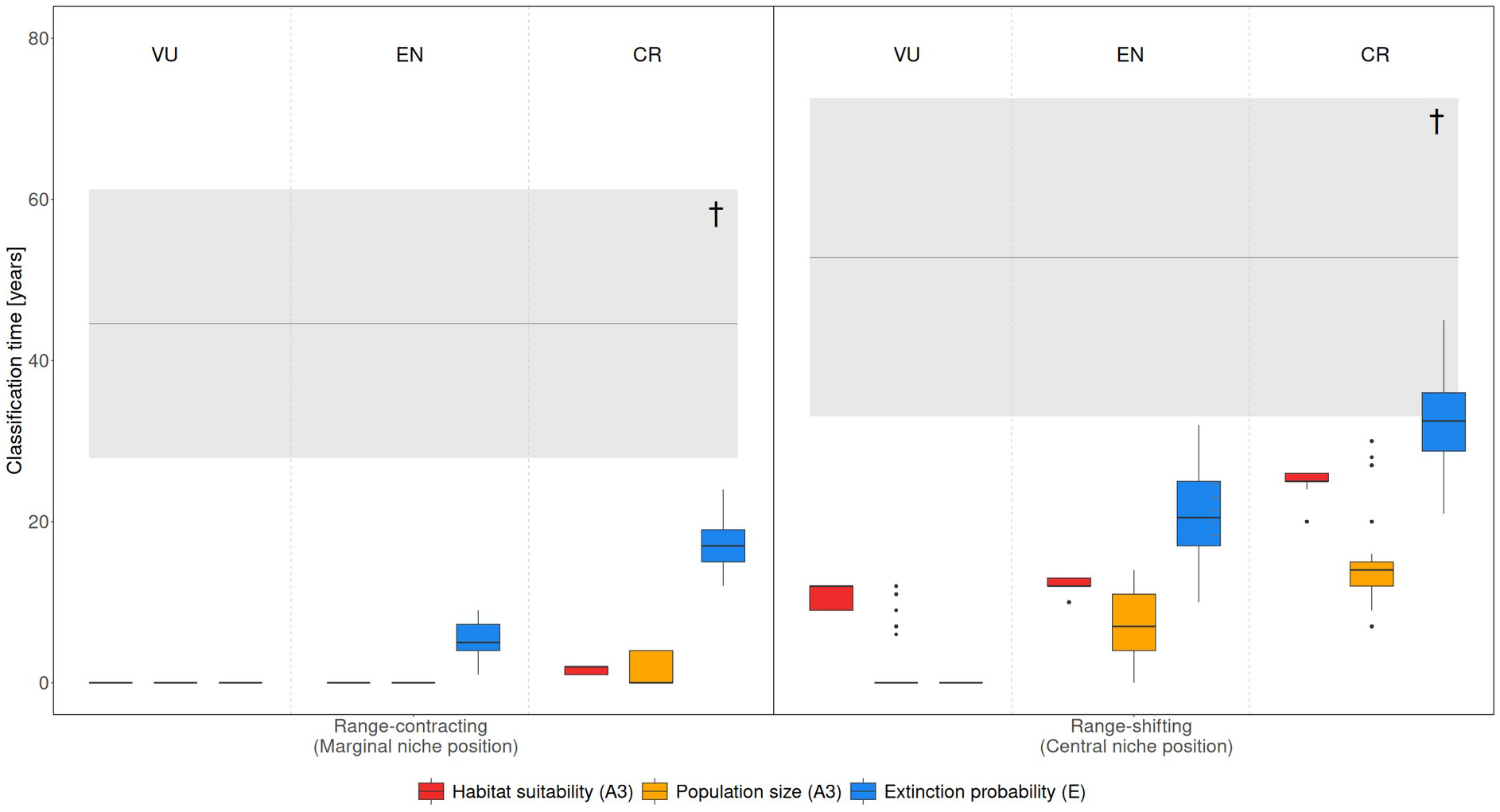
Classification times for listing species in different IUCN Red List threatened categories, compared to their extinction times. Results are shown separately for range-contracting species with marginal, cold-adapted niche position (left) vs. range-shifting species with central niche positions (right). Classification time refers to the years when species would first be classified into each of the three different threatened categories (Vulnerable, Endangered, Critically endangered). The boxplots show the classification times across simulated species, landscapes, and replicate runs using three approaches: true population size extracted from the simulation or SDM-derived habitat loss against criterion A3, and extinction probability calculated from the simulations against criterion E. Note that all species went extinct during the 90 years of climate change, thus qualifying for all threatened categories throughout their life span. The grey horizontal line shows the mean year of the extinction across species and the grey bands the 95% confidence interval (marked with †). Sampling size per box plot is n=240 (8 species x 3 replicate landscapes x 10 replicate runs).

For range-contracting species, the classification time did not differ between true population size and SDM-derived habitat loss (VU_pop_: 0±0 years; VU_habitat_: 0±0; EN_pop_: 0±0; EN_habitat_: 0±0; CR_pop_: 1.40±1.84; CR_habitat_: 1.74±0.44). By contrast, the classification of range-shifting species was substantially later when using SDM-derived habitat loss than when using true population size because range loss was underestimated by the SDMs (VU_pop_: 1.88±3.58 years; VU_habitat_: 10.71±1.39; EN_pop_: 6.67±4.52; EN_habitat_: 12.16±0.81; CR_pop_: 14.2±5.34; CR_habitat_: 25.07±1.21). This resulted in a very short warning time (i.e. time between the classification of a species and its extinction^6^) for range-shifting species when using SDM-based risk classification (Fig. 4 + Fig. S9). For range-shifting species that showed a full range shift (i.e., reaching the suitable habitat in the northern part of the landscape) due to high dispersal ability and fast growth, classification time between true population size and SDM-derived habitat loss did also not differ (Fig. S10). In a sensitivity analysis, we tested whether SDM-based predictions of range-shifting species could be improved by including simple dispersal assumptions but found no effect (Fig. S11). Other traits did not result in pronounced differences in classification times (Fig. S9). Interestingly, we found that for higher threatened categories (EN, CR) quantitative estimates of extinction probability and listing under criterion E resulted in considerably later classification time, and thus very short warning time, than information on population size and listing under criterion A3 (Fig. 4 + Fig. S9). The Red List guidelines indicate that criteria A-D are more precautionary than criterion E^8(p.17)^. Still, our results show that listing under these criteria can differ by up to 20 generations for our virtual species (Fig. 4; species were simulated with one generation per year).

## Discussion

Based on idealised simulated species and unbiased SDMs, we systematically evaluated the guidelines of the Red List for assessing species-level extinction risk under climate change. Our results revealed that the effectiveness of current Red List criteria strongly depend on expected range dynamics. SDM-based assessment provided sufficient warning time only for range-contracting species. For range-shifting species with time-lags in colonisation of newly suitable habitat, warning time was substantially shorter when using SDM-based assessment mainly because of the highly non-linear relationship between population size and SDM-predicted habitat loss. Other life-history traits had comparatively little effect on warning times. Further, model-based assessment based on probabilistic extinction estimates led to considerably shorter warning times than when based on predicted population sizes.

The strong effect of range dynamics stems from fundamental differences in how population size relates to predicted habitat loss. For range-contracting species, we found a convex relationship between population loss observed in the simulations and SDM-derived predictions of habitat loss. In these cases, species initially retained stable population sizes despite ongoing habitat loss while population sizes dropped very suddenly for higher rates of habitat loss. Such a convex relationship can be caused by a non-uniform distribution of individuals in the landscape, where sparsely populated edge habitats are lost first during range-contraction^21,34^, and by an extinction threshold, a critical amount of habitat beyond which population size declines steeply^23^. Crossing the threshold can initiate the extinction vortex, in which reinforcing processes such as inbreeding depression increase population decline, ultimately leading to species extinction^35^. Even though we did not consider species genetics or stage-structured populations in our simulation, we still observed an extinction threshold, which has been shown to be highly case specific and species dependent^23^. A convex shape of population decline over time was already reported for several species^36^, but research on the exact relationship between population size, habitat loss and extinction risk remains limited^20^. While the linear assumption in Red List guidelines is violated in these cases, our results suggest that SDM-based assessment still offers sufficient warning time for range-contracting species. Indeed, our results show that listing them based on SDM predictions could even lead to a slight overestimation of extinction risk for lower threatened categories (i.e., Vulnerable and Endangered) as the initial habitat loss was predicted to be higher than the actual population loss. It is important to note, however, that this did not result in misclassification but rather in slightly longer warning times, which could be beneficial for implementing effective conservation measures.

In contrast, range-shifting species exhibited a concave population size - habitat loss relationship indicating a considerable underestimation of range loss by SDM predictions, although the slope of the relationship differed between replicate landscapes. The concave curve likely reflects the inability of some species to fully track shifting climates due to dispersal limitations^37,38^ and time lags^39,40^. Certain landscape characteristics such as fragmentation, patch size and connectivity can enhance the dispersal ability of the species causing them to better track their suitable habitat and thereby influence the form of the curve^32^. The underestimation of range loss resulted in generally shorter warning time for SDM-based assessment of species in comparison to true population size. Because effective conservation measures often require several decades of advance warning, SDM-based assessment may not leave enough warning time to protect range-shifting species from extinction^29^. In contrast, SDM-based assessments did not lead to shorter warning time for fully range-shifting species with high dispersal ability and fast growth, highlighting the relevance of explicitly considering dispersal and demographic processes in climate impact assessments^17^. Stronger dispersal ability is expected to improve climate tracking and the survival of range-shifting species under climate change^32,41^. Interestingly, we only found such influence of dispersal ability on the classification time for one range-shifting species and found delayed warning for all other range-shifting species. As inclusion of simple dispersal assumptions did not change these results, our study suggests that SDMs will not be adequate for assessing extinction risk in range-shifting species threatened with climate change. For these species, spatially explicit population models are likely more adequate.

Our results further revealed that Red Listing based on probabilistic extinction estimates under criterion E led to delayed classifications compared to criterion A3 and insufficient warning times for all species, particularly those most at risk. This aligns with previous findings that criterion E often results in less precautionary extinction risk status in comparison to other criteria^42^. In our study, the short warning time resulted from steep, sigmoidal extinction risk curves, which led to rapid population collapse and abrupt extinction of species. Such curve shapes have been shown to realistically describe species’ survival probabilities under changing environmental conditions^43,44^. In stochastic population models, sigmoidal extinction curves typically emerge when full parameter knowledge during the model building process is available^45^. With higher parameter uncertainty the extinction risk curves are expected to become more shallow^45^, which consequently influences the classification time in the Red List. Interestingly, a previous study^29^ reported that criterion A leaves less warning time in comparison to listing against criterion E when using a coupled SDM-population model approach with a real species. It is thus unclear whether a higher parameter uncertainty in the model building process for criterion E might leave more warning time or may introduce additional biases. As a more robust approach for species threatened by climate change, especially those undergoing range shifts, we recommend assessing extinction risk based on predicted abundances or expected minimum abundance (EMA) from SEPMs, as assessment using predicted abundances (i.e. population size) lead to longer warning times. Our results suggest that assessing extinction risk based on these projections provides a more reliable basis for classification than relying on quantitative extinction probabilities alone.

Our simulations were idealised, with full knowledge of the true population and range dynamics, perfect (i.e., error-free and unbiased) occurrence data to fit SDMs, and perfect knowledge and information on the environmental drivers, which allowed the estimation of unbiased SDMs. In practice, missing data, sampling biases, and missing knowledge about species ecology and dispersal as well as environmental and demographic stochasticity could introduce additional uncertainties in the models and thus in the assessment process^46–48^. Here, we want to echo the Red List guidelines in stressing transparency regarding uncertainties in the assessment process to minimize their effect^8(pp.23-25)^. Additionally, our climate change scenario was simplistic with only temperature changing over time. Simulating more driver variables and more realistic climate change scenarios, which include extreme events and more realistic fluctuations in temperature and precipitation, could lead to more diverse responses of species. Furthermore, more detailed species characteristics, such as stage-structured populations, stochastic movement dispersal and density-dependent traits could also influence the responses of the species and thereby the assessment process. Nevertheless, our results revealed that even under idealised conditions, current Red List criteria are highly sensitive to species range dynamics under climate change.

To improve assessment, we recommend including a screening step into model-based Red List assessments. In this initial screening, SDMs can serve to assess whether a species is likely to experience exclusive range contraction or whether suitable habitat is likely to shift geographically. If range-shifting is likely, we strongly recommend using spatially explicit population models and assess extinction risk based on predicted abundance or the expected minimum abundance as used in population viability analyses^47^, rather than extinction probabilities. While adjusting thresholds for extinction probabilities under criterion E or the time frames in general may improve assessments in some cases, further research is needed to define thresholds and time frames that offer both realism and sufficient warning time. Applying dispersal buffers did not improve Red List classification for range-shifting species in our examples, and we suggest that future studies could test the applicability of more sophisticated and temporally finer resolved dispersal models in this context. For range-contracting species, SDMs may be appropriate to assess the extinction risk of species. Still, the assumption of a linear relationship between population size and habitat size should be treated with caution as our results show that sudden population reductions can cause delayed warning by SDMs. Furthermore, the known limitations of SDMs should be kept in mind^18^ and the assessment based on SDMs should follow best practices^49,50^ to avoid wrong classifications. Here, we want to stress again that our SDMs represented the ideal case with perfectly known and unbiased data while in reality species ecology, dispersal and observer behaviour could bias SDMs. To further guide the novelization of the Red List under climate change, we recommend empirically testing our results for a variety of species for which adequate range loss and population loss data are available.

## Methods

We used the RangeShifter framework^33,51,52^ to simulate population dynamics of sixteen virtual species under equilibrium conditions and under climate change in three different artificial landscapes. Species differed in four traits: niche position (central vs. marginal), niche width (narrow vs. wide), growth rate (slow vs. fast) and dispersal (short vs. long), leading to 16 (2^4^) different species to cover all trait combinations. Niches were defined by two environmental variables that could be interpreted as temperature and precipitation. We aimed to simulate a representative set of virtual species spanning a wide range of different traits and characteristics as well as extinction dynamics of these species to assess their effect on the classification in the IUCN Red List. Species occurrence data and climate data were extracted from the simulation at equilibrium (after 100 years of spinup) before climate change set in. Based on these data we fitted species distribution models (SDMs) and predicted habitat suitability under climate change. From the simulations, we observed population size across the entire time frame of 90 years of climate change and estimated extinction probability per year. We ran simulations for 90 years of climate change to ensure the extinction of all species. From the SDMs, we derived the predicted habitat suitability values for each of these 90 years and compared these to true population size. Last, we applied the Red List criteria to each of these three metrics to classify species into different threatened categories.

### Virtual species

In our simulation approach, we first modelled the population dynamics of the 16 virtual species in the three artificial landscapes under climate change. The three artificial landscapes are stochastic replicates obtained using a neutral landscape model^53^. The artificial landscape consisted of two environmental variables (which could be interpreted as temperature and precipitation) with a spatial extent of 512 x 512 cells at 1000 m resolution. The environmental layers were generated using the package “NLMR”^54^. Temperature was represented by a latitudinal gradient with spatially auto-correlated noise. For this, the arithmetic mean of a landscape with a linear gradient and a landscape built with midpoint displacement with a Hurst exponent of 0.96 was taken. To create spatially auto-correlated precipitation patterns, we generated a landscape using midpoint displacement with a Hurst exponent of 0.75. Landscape values ranged between 0 and 1. We derived three replicate landscapes using the same settings (Fig. S12).

The virtual species niche was modelled using the package “virtualspecies”^55^. First, Gaussian-shaped response curves for both environmental variables were simulated. The mean of the response curve for temperature was adapted to the respective niche position (marginal: 0.27, central: 0.5), the mean of the precipitation response curve was set to 0.5. To model different niche breadths, the standard deviation of both response curves was adapted based on the respective niche breadth (narrow: 0.045, wide: 0.055). The response curves of the simulated niches are included in the appendix (Fig. S13). We then used the product of both response curves to map the virtual species niche in the artificial landscapes. The different niche positions (i.e., marginal and central) were explicitly selected to model different range dynamics to test their influence on the extinction of the species and thus their classification in the IUCN Red List. A marginal, cold-adapted niche position caused range contraction under climate change and extinction, while a central niche position led to range-shifting prior to range contraction and extinction. We decided to only model cold-adapted marginal niches in the north, as a warm-adapted marginal niche position in the south would have resulted in the same range-shifting pattern as the central niche position but with a longer time period for the range-shift.

To simulate climate change, temperature was linearly increased over 90 years (Fig. S14), while precipitation remained static. We selected a time frame of 90 years to model a continuous increase of temperature that would result in no suitable habitat being left at the end of climate change, ensuring the extinction of all species. To the linear temperature increase we also included a temporal autocorrelated noise. For this we used an ARIMA model^56^ of the order 0,0,1 and added it to the linear model using the following equations:

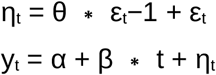

Where, η_t_ represents the time series obtained from the ARIMA model with θ set to 0.5 and ε representing the residuals at the respective time points. This time series was then added to a linear model with α set to 0.5, β set to 0.6, and t representing the respective year. The time series was rescaled between 0 and 0.9 and added to the temperature value of every cell in the artificial landscape. Values that were larger than 1 were reduced to 1 to stay within the range of 0 to 1. We did not aim to mimic realistic climate change projections but modeled climate change in a way to force the species to extinction within a reasonable time frame. Precipitation remained static under climate change and extreme events were not considered to reduce complexity.

To model the population dynamics of the virtual species, we used the spatially explicit and individual-based modelling platform RangeShifter^33,51^, operated via the package “RangeShiftR”^52^. We implemented a cell-based model using the previously created maps of the virtual species niche with the resolution set to 1000 m and the carrying capacity set to 0.05 ind/ha. We modelled the virtual species as an asexual species with non-overlapping generations meaning that each individual only lived for one year. The competition coefficient was set to 1 (i.e., no over- or undercompensation) and the value for the maximum growth rate was adapted according to the growth rate (slow: 3, fast: 5). After reproduction the adults died and the offspring dispersed. Dispersal is modelled explicitly in three phases in RangeShifter: emigration, transfer, and settlement. The emigration probability for each individual was set to 0.4 and was density-independent. The transfer was modelled using a negative exponential dispersal kernel. For short-distance dispersal, the mean dispersal distance was set to 5000 m. For long-distance dispersal, a double exponential dispersal kernel was used to include rare long-distance dispersal events. The double kernel is composed of two negative exponential kernels with different means and probabilities of occurrence. The first (shorter) kernel occurred with a probability of 95% and had a mean dispersal distance of 15 000 m, the second (longer) kernel occurred with a probability of 5% and had a mean dispersal distance of 250 000 m. If the arrival point of an individual went beyond the boundaries of the landscape or within the natal cell the dispersal process was repeated. The individual settled in the arrival cell if the cell contained suitable habitat. If the arrival cell was not suitable, the individual was displaced to one of the eight neighbouring cells, if one of them was suitable, or else died. If a cell became unsuitable due to climate change, all individuals inhabiting that cell died immediately.

Trait values were selected based on two criteria: all species had to achieve a stable population size after the spin-up period of simulation, while the simulation still had to be computationally feasible and the differences between the trait values had to be large enough to show different effects. We used realistic species (i.e., small rodents and birds) as guidance for the determination of the growth rate and dispersal distances.

The simulation was initialised using the map with equilibrium climatic conditions and individuals were distributed in all suitable cells at half-carrying capacity. The simulation ran for 100 spin-up years to remove any initialisation effect. After the spin-up period, the landscape was changed yearly to represent the virtual species niche shifting northwards in response to climate change. In the following, we refer to year 0 as the last year of equilibrium (after spinup) and year 1-90 represent the years under climate change. Each of the 16 species was simulated with 100 replicate runs for each of the three replicate artificial landscapes.

From the output of each simulation, we obtained the population size of the species for each year and replicate run. To calculate the extinction probability for each year, we divided the number of replicate runs without a viable population (i.e., no individual was left in the landscape) of that year by the total number of replicate runs^57^.

### Species distribution models

At the end of the spin-up period, before climate change set in, we extracted species presence and absence information (Fig. S2) as well as climate variables from the simulation to fit SDMs. To account for variation between the replicate runs, we randomly selected ten replicate runs from the simulation and fitted separate SDMs to each. This resulted in 480 SDMs (16 species x 3 landscape replicates x 10 replicate runs). For the species data we assumed perfect sampling with no error in detection. Data points were partially thinned by selecting every second cell to avoid spatial autocorrelation using the package “dismo”^58^. Based on the recommendations of the Red List^8(p.108)^ we used three distinct modelling algorithms (Generalised Linear Model (GLM), Random Forest and MaxEnt) and further calculated an ensemble model based on the arithmetic mean. For the GLM, linear and quadratic predictors were used with a binomial error distribution. For Random Forest, we fitted 1000 trees with a minimum terminal node size of 20 to fit comparably smooth response surfaces. In the MaxEnt algorithm, we used the feature classes linear, quadratic and hinge.

The probability of occurrence was predicted for each of the 90 years under climate change using the ensemble model. The predictive performance of the model algorithms and the ensemble model was evaluated using the 99 replicate runs that were not used for fitting the SDM. As performance measures we selected the AUC (area under the ROC curve) and TSS (true skill statistics)^16^. The calculated performance measures were averaged over all 99 replicate runs using the arithmetic mean. For the calibration and evaluation of the SDMs we used the following packages: “randomForest”^59^, “maxnet”^60^, “gbm”^61^, “PresenceAbsence”^62^ and “lhs”^63^.

For testing the effect of dispersal assumptions in SDMs on the assessment in the Red List, we first binarized the predictions of each species and landscape replicate using the maxTSS threshold^64^. Around the known presences we created a buffer with the size of the estimated dispersal distance of an individual in the respective time frame and set the predicted habitat suitability outside the buffer to 0 following the recommendation of the Red List guidelines^8(p.110)^. Buffer sizes were determined by the empirical dispersal distances observed from the RangeShifter simulations. For this, we extracted the dispersal distances of all emigrated individuals for year 0 from the simulations, and calculated the arithmetic mean of dispersal distance.

### Red List classification

To assess Red Listing of species threatened by climate, we compared the assessment of the species against two different Red List criteria using three metrics. For the criterion A3, we used true population size observed from the RangeShifter simulations and the sum of the SDM-derived habitat suitability values. For the criterion E, we used the extinction probability for every species obtained from the RangeShifter simulations. For the population size, we only used the population size from the ten replicate runs that were selected for fitting the SDMs. To calculate the sum of the habitat suitability values from the SDM predictions we followed the recommendations from the Red List guidelines^8(p.99)^. We first removed cells with values below a certain threshold value, for which we used the maxTSS^64^. We then summed up the predicted habitat suitability of the remaining cells and for every year. For the classification into the different categories of the Red List, different thresholds and time frames apply for the different criteria (see Tab. 1).

For each species, we determined the time of first classification into the three different threatened categories using all three metrics (i.e., population size, habitat suitability and extinction probability). As the Red List criteria are tied to specific time frames, we to tested for every consecutive year if the species would fulfill the Red List criteria in the following years (see Table 1 for considered time spans). If the criterion was fulfilled, then this particular year was noted as the time of classification for that criterion. If the criterion was not fulfilled, we repeated the analyses for the next consecutive year until we found the year in which the species would first be classified according to the criterion under question. As all species continuously experienced population and habitat loss under climate change until their extinction, each species qualified for all threatened categories throughout their life span (within the 90 years of simulated climate change).

**Table 1:** Thresholds and time frames used to classify species into threatened categories, based on the Red List Criteria^8^.

| Criterion | Red List Threatened Categories |  |  |
| --- | --- | --- | --- |
|  | Critically Endangered | Endangered | Vulnerable |
| A3: future population size reduction / future reduction of sum of SDM-derived habitat suitability | $\geq 80\%$ in 10 years | $\geq 50\%$ in 10 years | $\geq 30\%$ in 10 years |
| E: probability of extinction | $\geq 50\%$ in 10 years | $\geq 20\%$ in 20 years | $\geq 10\%$ in 100 years |

In a sensitivity analysis, the assessment of classification times based on SDM-derived habitat suitability values was repeated for more realistic dispersal assumptions using the buffer approach described earlier.

### Statistical analysis

To quantify the relative importance of the different traits for explaining differences in the population size - habitat loss relationship, we fitted a generalised mixed-model ANCOVA using an ordered beta regression implemented by the package “ordbetareg”^65,66^. This package is a front-end to the package “brms”^67^. We selected an ordered beta regression since our response variable (simulated “true” populated size) was continuous with a zero and one inflation. As predictors, we considered habitat loss as continuous variable, the four traits (niche position, niche width, dispersal distance and growth rate) as categorical variables with two levels each, and their pairwise interactions with habitat loss. We further controlled for the landscapes by including them as random intercepts. The best model was selected based on the difference in expected log predictive density. Posterior distributions were estimated using Markov chain Monte Carlo (MCMC) sampling with 4 chains and a delta value of 0.95. Each chain had a length of 6000 iterations, with the first 2000 being used as a warm-up. For the prior, the default was selected.

For the study R 4.2.2^68^ was used. In addition to the already mentioned packages the following were used for data preparation and visualization: “scales”^69^, “dplyr”^70^, “ggplot2”^71^, “terra”^72^, “gridExtra”^73^, “tibble”^74^, “RColorBrewer”^75^, “doParallel”^76^, “foreach”^77^, “data.table”^78^, “ggtext”^79^.

## Code availability

The codes for this study are available at https://doi.org/10.5281/zenodo.19236947

## Author Contributions

RK and DZ conceptualised the study. RK ran all analyses and led the writing. All authors contributed to revising the text.

## Acknowledgments

This study was supported by the German Research Foundation DFG (grant no. 518316503). S.A.F. gratefully acknowledges the support of iDiv funded by the German Research Foundation (DFG– _FZT 118, 202548816) and of the Leibniz Competition by the Leibniz Association (P52/2017). We thank Cara Gallagher for helpful discussions during early phases of the work.

