## Supplement for "Red List criteria underestimate climate-related extinction risk of range-shifting species"

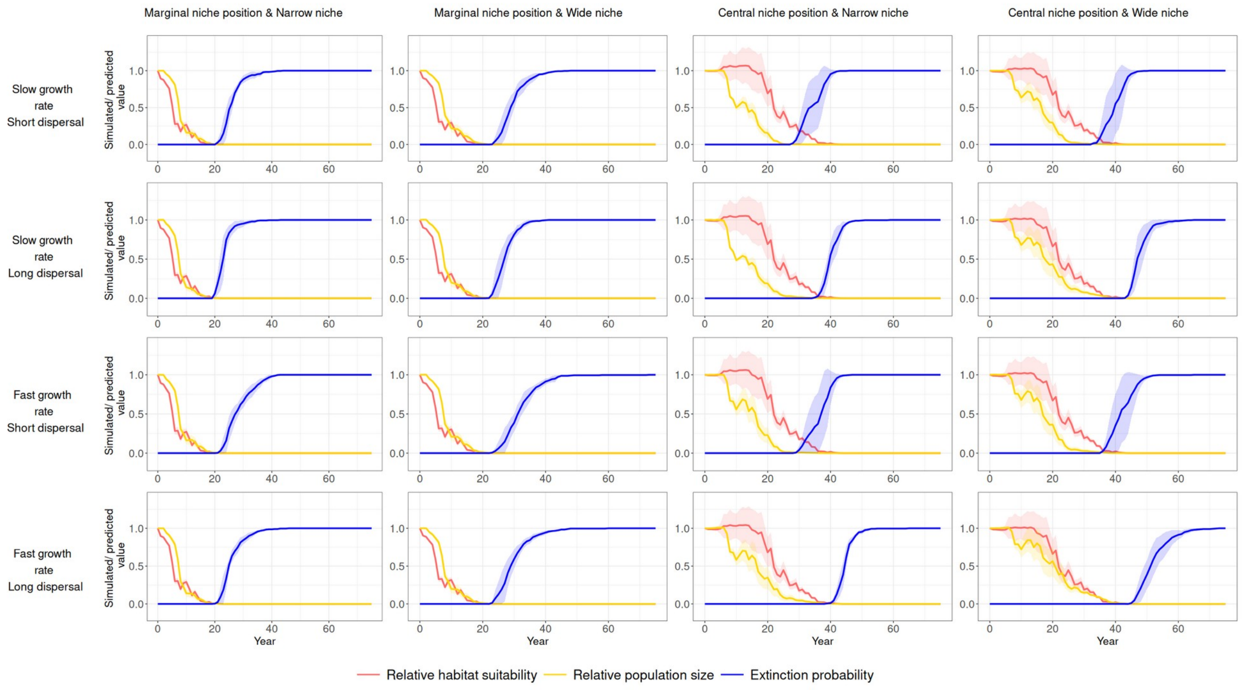


Figure S1: Time trajectory of relative population size, SDM-derived habitat suitability and extinction probability for the 16 different species. Relative population size is obtained from the RangeShifter simulations as „true“ metric for the virtual species. Extinction probability is calculated from the RangeShifter simulations. All metrics are averaged across ten replicate runs. Confidence bands display the differences across three landscape replicates and ten replicate runs. Species with marginal niche positions showed range-contracting under climate change. Species with central niche positions showed range-shifting under climate change.

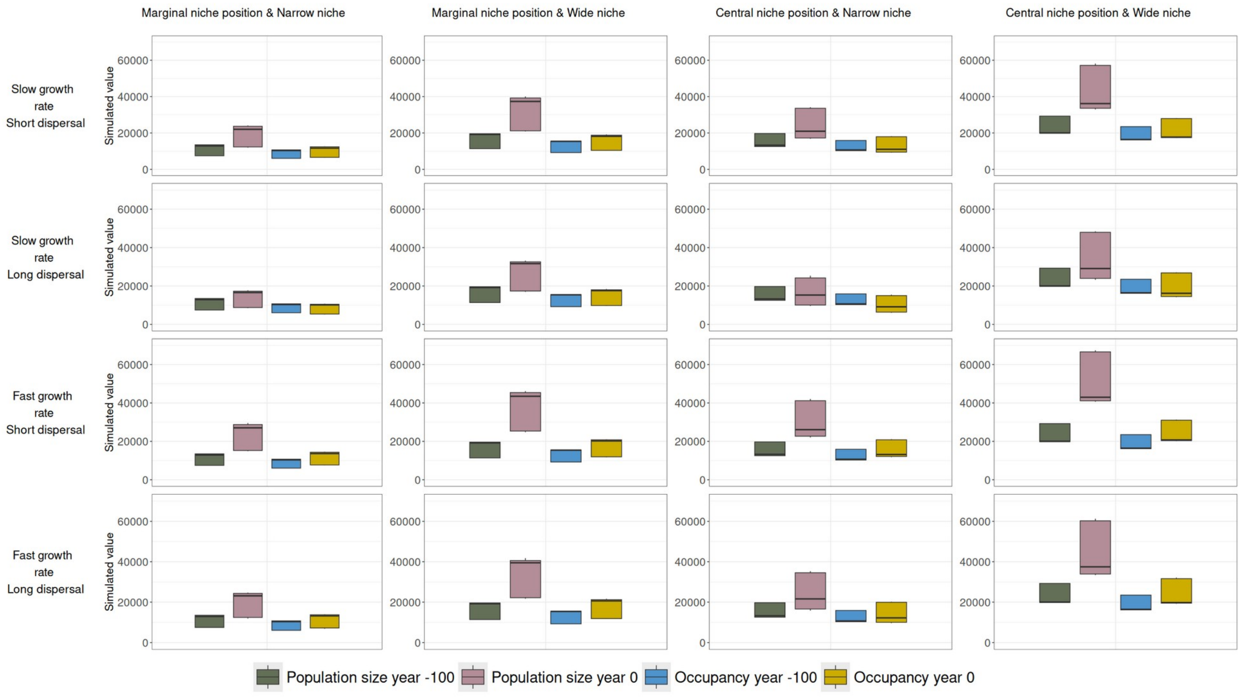


Figure S2: Boxplots showing the population size and occupancy (number of occupied cells, i.e. range size) at the start of the spinup of the RangeShifter simulation (Year -100) and at equilibrium after 100 years of spinup (before climate change sets in) for the 16 different species. Sampling size per boxplot is n=30 (3 replicate landscapes x 10 replicate runs).

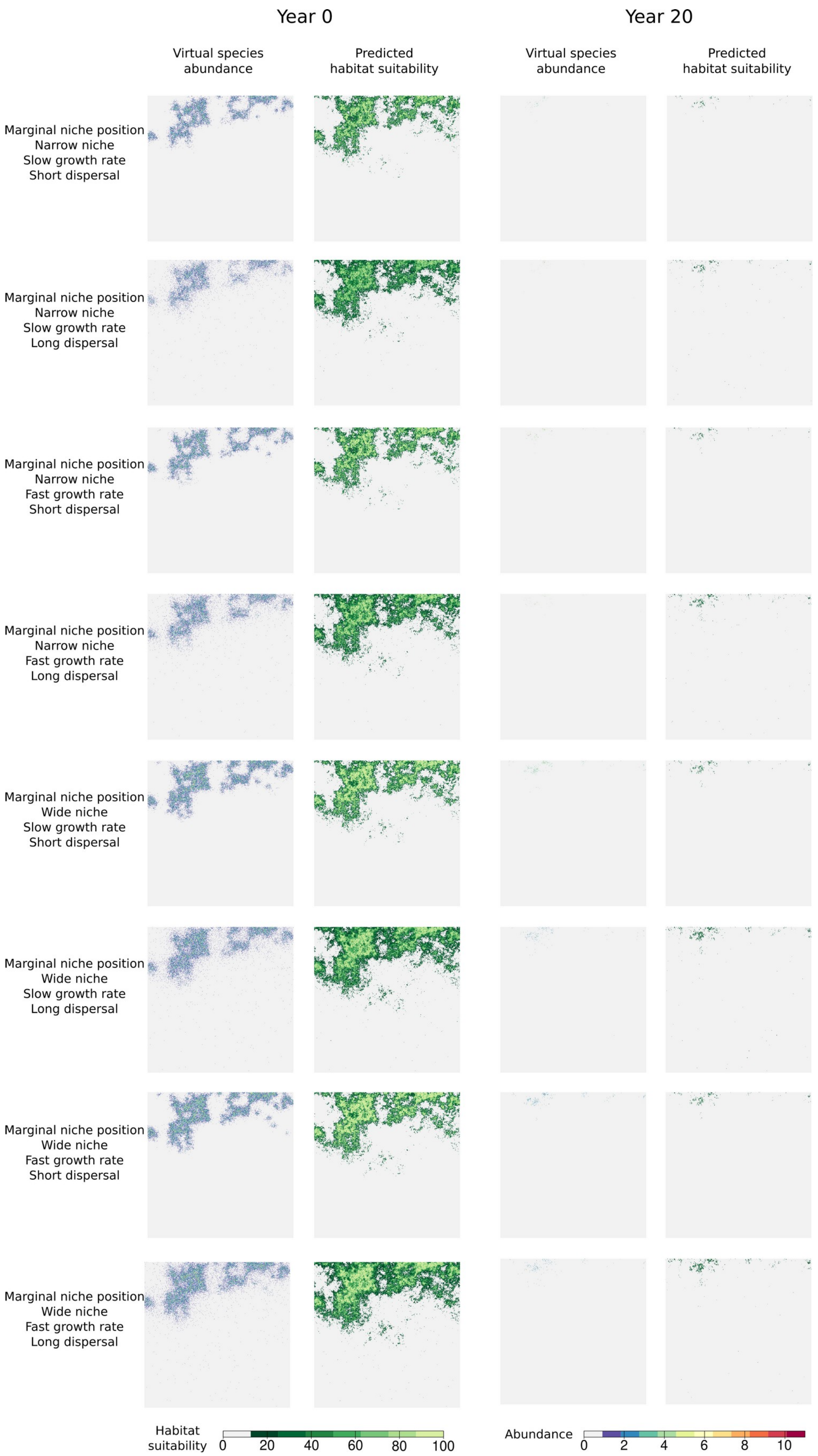


Figure S3: Maps of simulated „true“ species abundance and maps of habitat suitability predicted by SDMs for species with a marginal, col-adapted niche position. Results are shown for one replicate run under equilibrium conditions (year 0) and under climate change (year 20; shortly before extinction, cf. Fig. S1) in one example landscape. Species with marginal niche positions showed range-contracting under climate change.

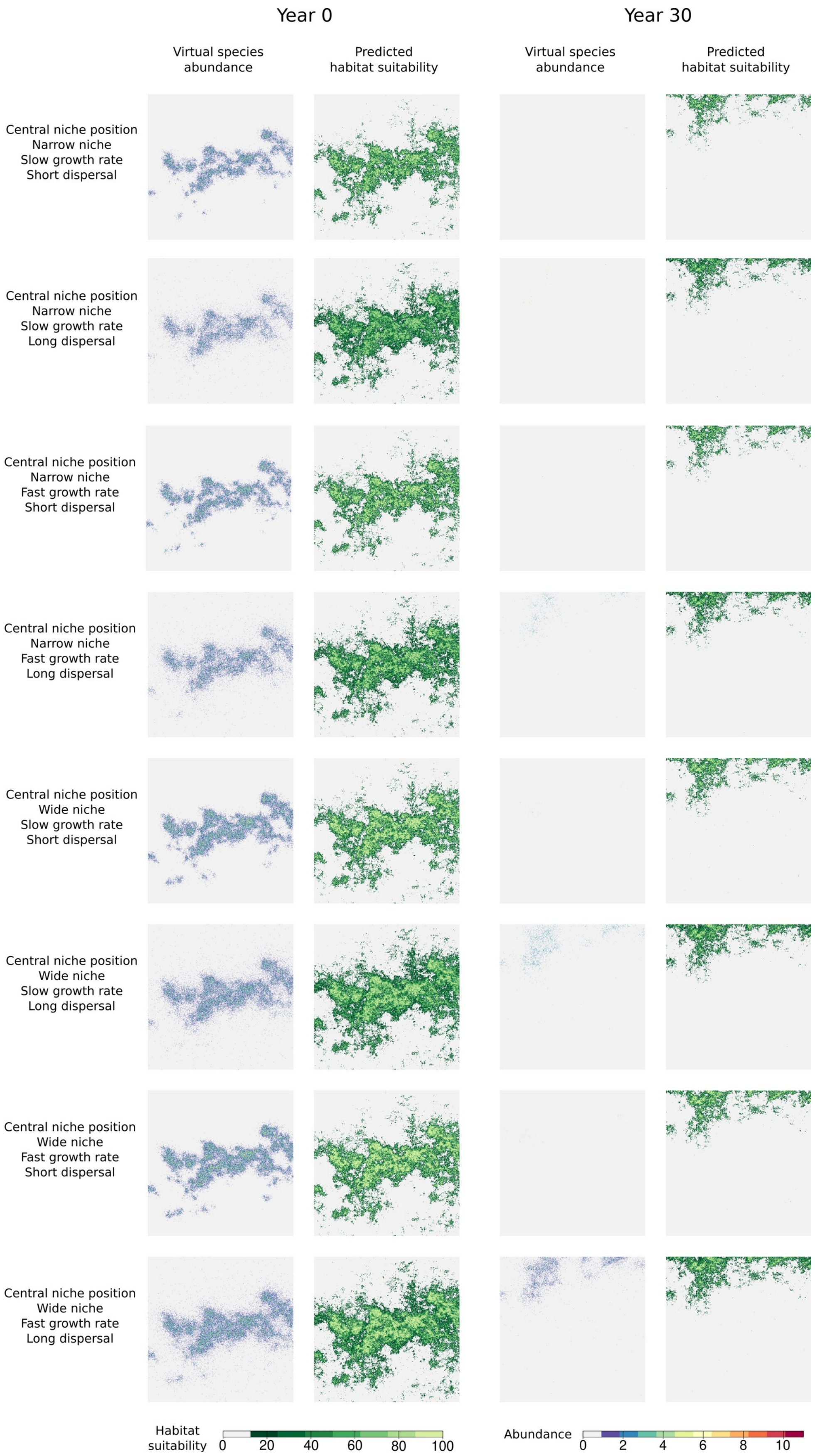


Figure S4: Maps of simulated „true“ species abundance and maps of habitat suitability predicted by SDMs for species with a central niche position. Results are shown for one replicate run under equilibrium conditions (year 0) and under climate change (year 30; shortly before extinction, cf. Fig. S1) in one example landscape. Species with central niche positions showed range-shifting under climate change.

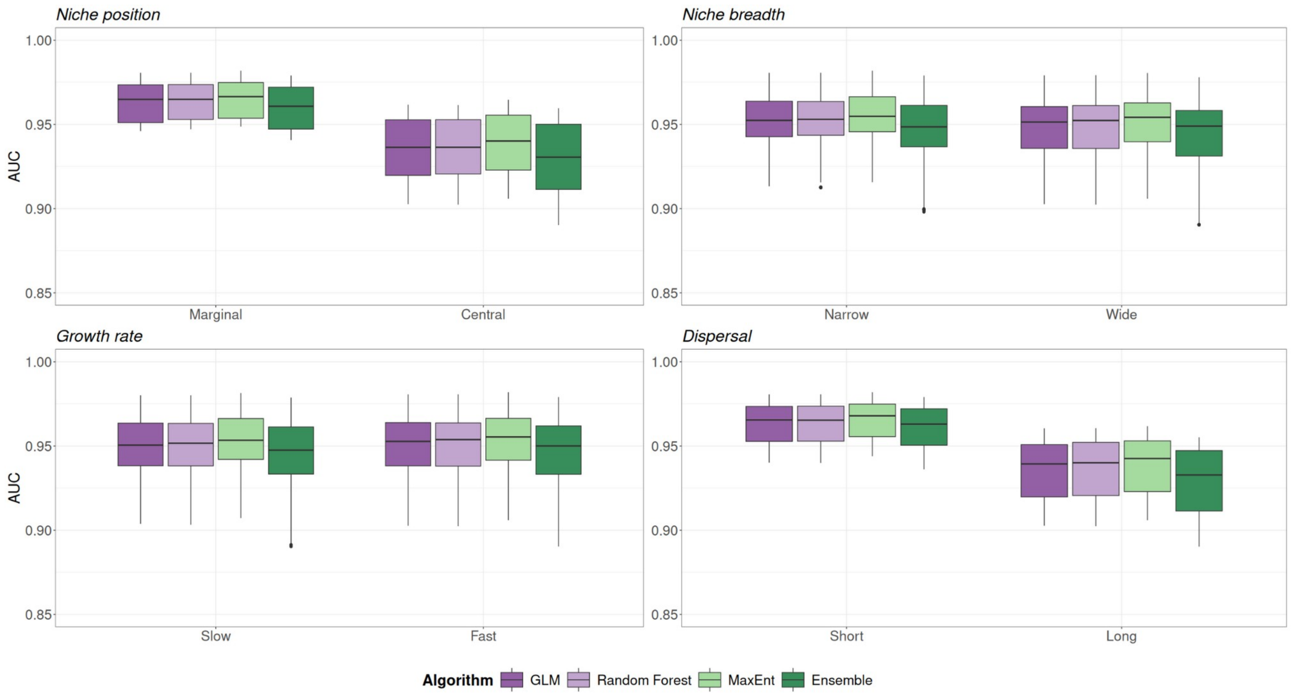


Figure S5: Boxplots of the AUC values for the different SDM algorithms and the ensemble model by traits and trait levels for all species, landscape replicates and replicate runs. SDMs were evaluated prior to climate change. Sampling size per box plot is n=240 (8 species x 3 replicate landscapes x 10 replicate runs).

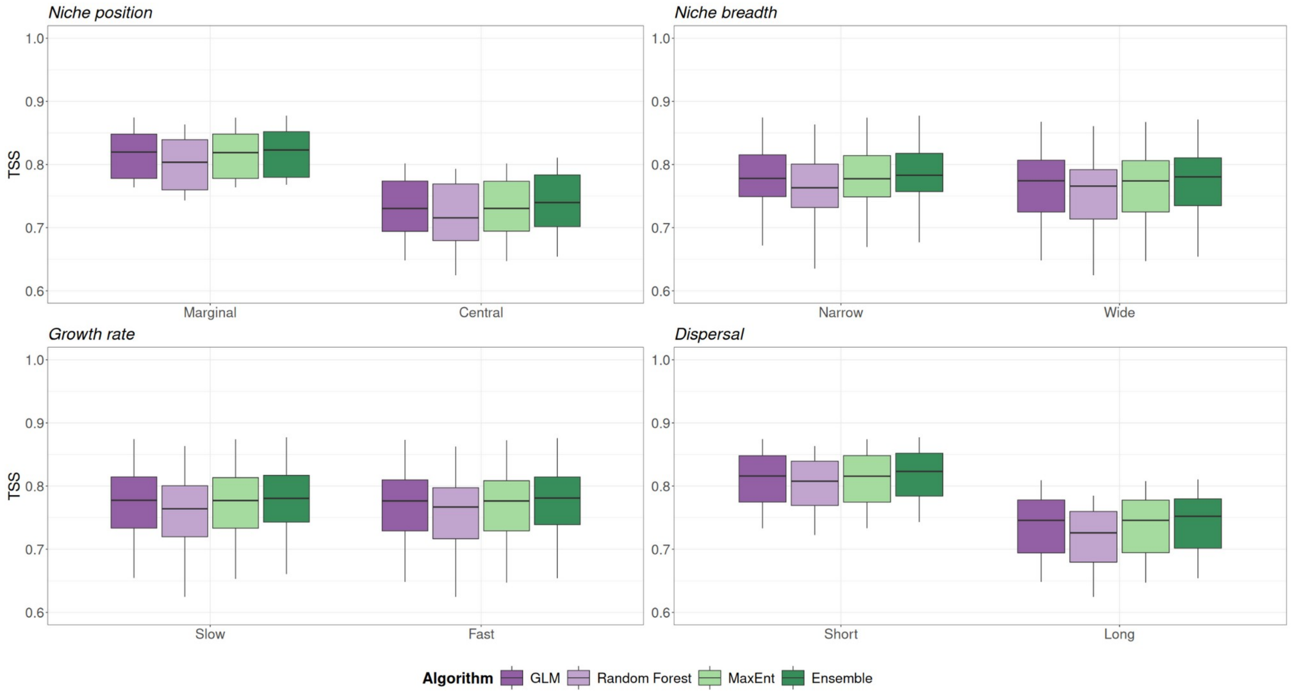


Figure S6: Boxplots of the TSS values for the different SDM algorithms and the ensemble model by traits and trait levels for all species, landscape replicates and replicate runs. SDMs were evaluated prior to climate change. Sampling size per box plot is n=240 (8 species x 3 replicate landscapes x 10 replicate runs).

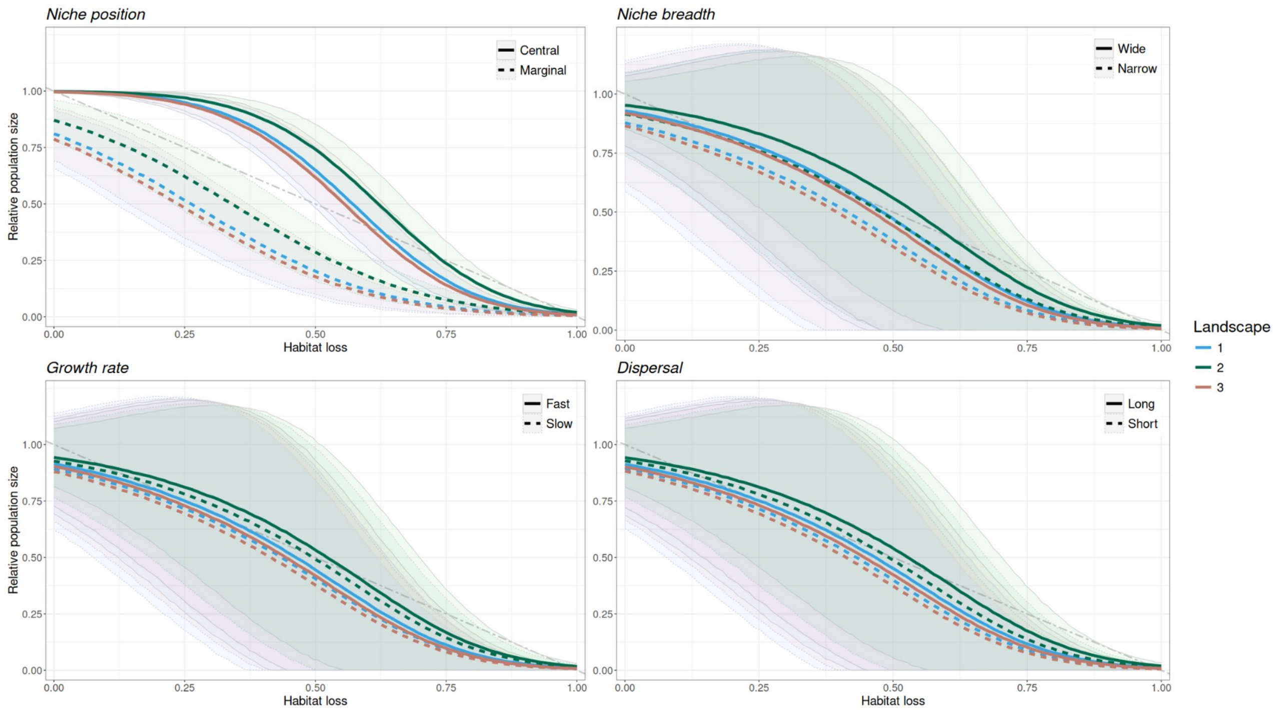


Figure S7: Relationship between relative population size and habitat loss for all simulated species. Relative population size was obtained from the simulation (observed true data for virtual species) and habitat loss was predicted by the SDMs. The dot-dash grey line shows the assumption of a linear relationship between population size and habitat loss as proposed by the Red List guidelines. The relationships are displayed using an ordered beta regression model, which was fitted using the “true” population size values obtained from the simulations as response variable and habitat loss, the 16 trait combinations across the 10 selected replicate runs as predictor variables, and landscape as a random effect. Lines indicate the mean model predictions and the transparent bands the 95% confidence interval. Species with marginal niche positions showed range-contracting under climate change. Species with central niche positions showed range-shifting under climate change.

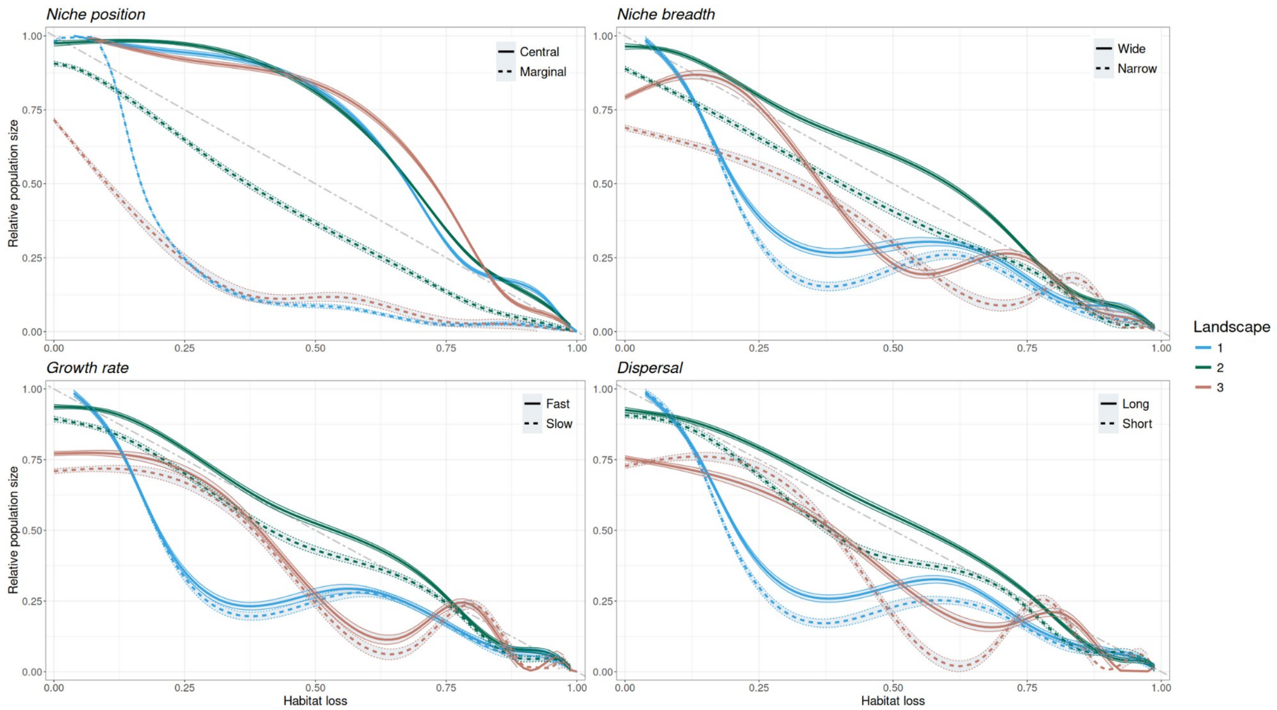


Figure S8: Relationship between relative population size and habitat loss visualised with loess smoothing. Relative population size was obtained from the simulation (observed true data for virtual species) and habitat loss was predicted by the SDMs. The dot-dash grey line shows the assumption of a linear relationship between population size and habitat loss as proposed by the Red List Guidelines. The relationships are displayed using a loess smoother on the simulation outcomes for each trait level and landscape separately. Lines indicate the mean predictions and the transparent bands the 95% confidence interval. Species with marginal niche positions showed range-contracting under climate change. Species with central niche positions showed range-shifting under climate change.

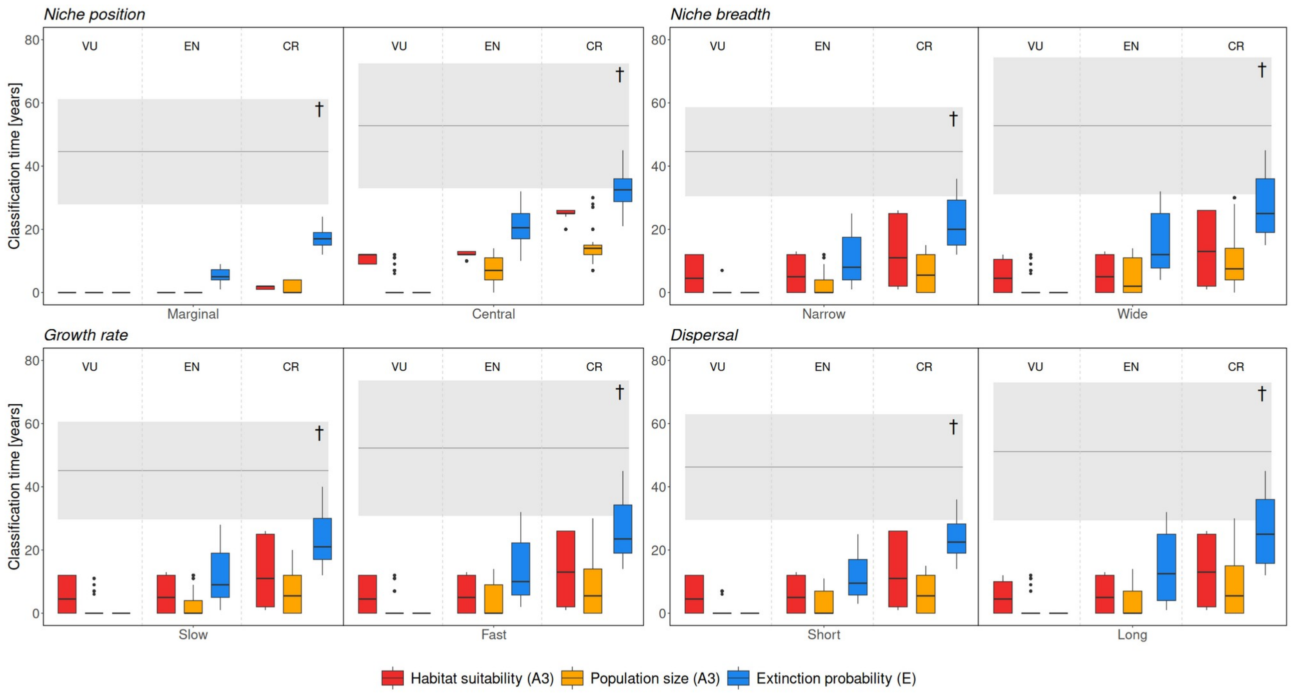


Figure S9: Classification times for listing the different species in the IUCN Red List threatened categories. Classification time refers to the years when species would first be classified into each of the three different threatened categories (Vulnerable, Endangered, Critically endangered). Classification time refers to the years when species would first be classified into each of the three different threatened categories (Vulnerable, Endangered, Critically endangered). The boxplots show the classification times across simulated species, landscapes, and replicate runs using three approaches: true population size extracted from the simulation or SDM-derived habitat loss against criterion A3, and extinction probability calculated from the simulations against criterion E. Note that all species went extinct during the 90 years of climate change, thus qualifying for all threatened categories throughout their life span. The grey horizontal line shows the mean year of the extinction across species and the transparent bands the 95% confidence interval (marked with †). Species with marginal niche positions showed range-contracting under climate change. Species with central niche positions showed range-shifting under climate change. Sampling size per box plot is n=240 (8 species x 3 replicate landscapes x 10 replicate runs).

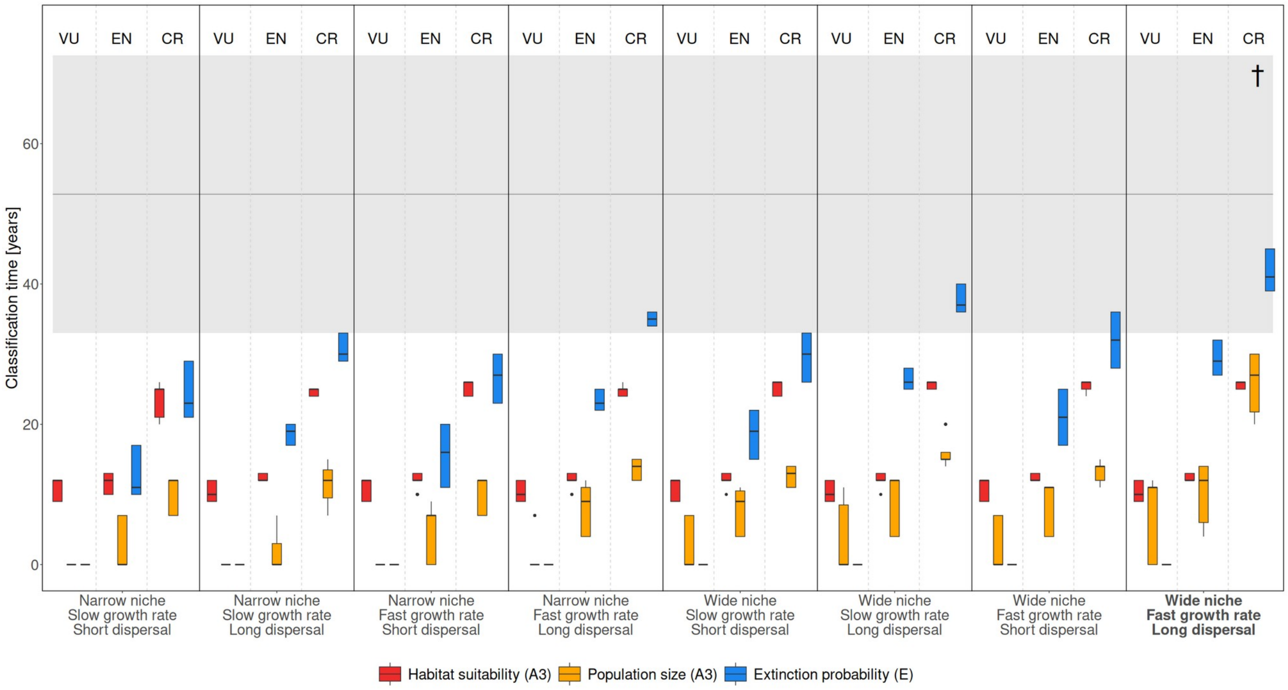


Figure S10: Classification times for listing the species in the IUCN Red List threatened categories for all eight range-shifting species separately. Classification time refers to the years when species would first be classified into each of the three different threatened categories (Vulnerable, Endangered, Critically endangered). The boxplots show the classification times across simulated species, landscapes, and replicate runs using three approaches: true population size extracted from the simulation or SDM-derived habitat loss against criterion A3, and extinction probability calculated from the simulations against criterion E. Note that all range-shifting species showed a range-shift, but some species were dispersal limited (i.e. not able to reach the suitable habitat in the northern part of the landscape). Species showing a full range shift and no dispersal limitation (i.e. reaching the suitable habitat in the northern part of the landscape) are marked in bold. Note that all species went extinct during the 90 years of climate change, thus qualifying for all threatened categories throughout their life span. The grey horizontal line shows the mean year of the extinction across species and the transparent bands the 95% confidence interval (marked with †). Species with marginal niche positions showed range-contracting under climate change. Species with central niche positions showed range-shifting under climate change. Sampling size per box plot is n=30 (3 replicate landscapes x 10 replicate runs).

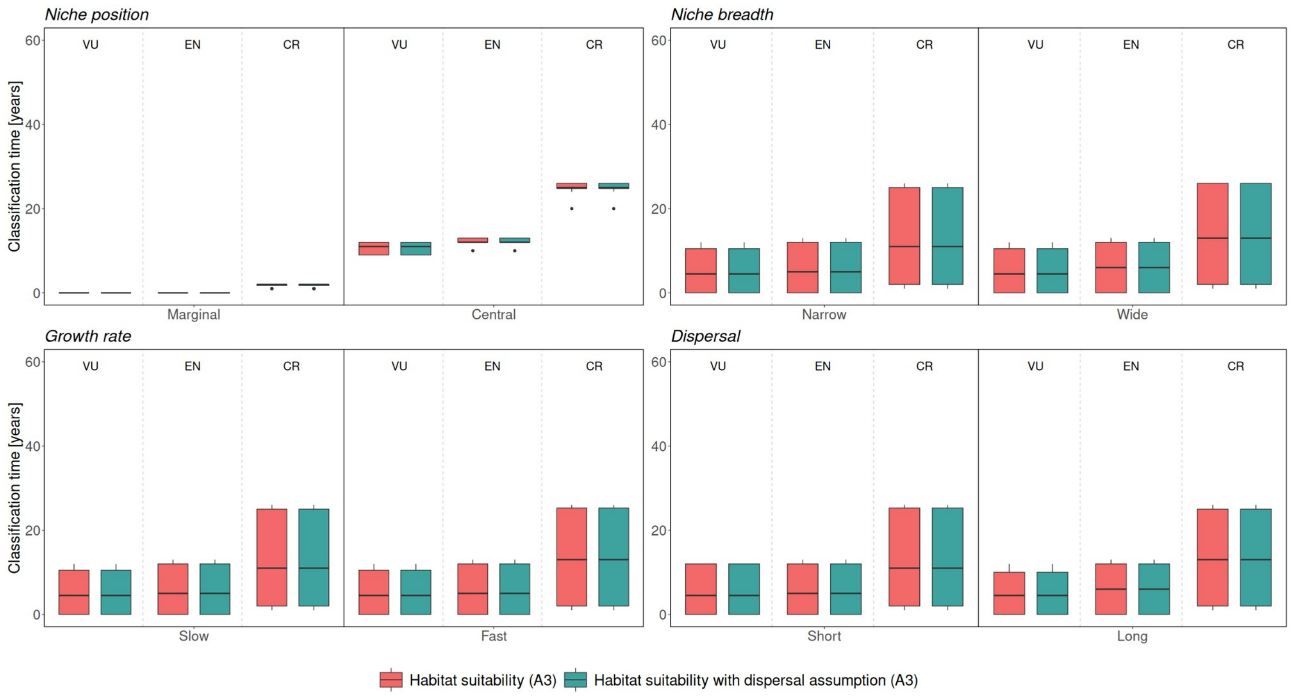


Figure S11: Results of the sensitivity analysis for the effect of dispersal assumptions on SDM-based risk classification. Displayed are the classification time points for listing the different species in the IUCN Red List threatened categories (Vulnerable, Endangered, Critically Endangered). Classification time refers to the years when species would first be classified into the three threatened categories using SDM-derived habitat loss and corrected SDM-derived habitat loss using dispersal assumptions under criterion A3. Dispersal assumptions are based on the empirical dispersal distances observed from the RangeShifter simulations. Species with marginal niche positions showed range-contracting under climate change. Species with central niche positions showed range-shifting under climate change. Sampling size per box plot is n=240 (8 species x 3 replicate landscapes x 10 replicate runs).

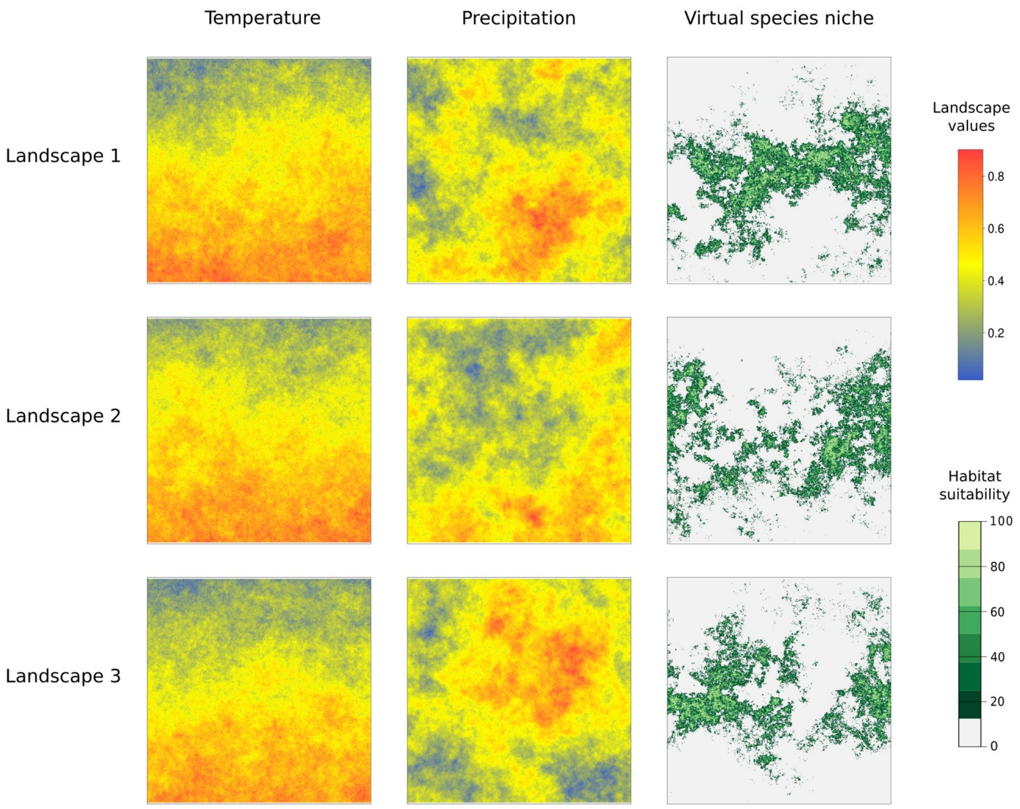


Figure S12: The three artificial landscapes of temperature and precipitation and the corresponding habitat suitability maps of the virtual species niche from a species with a central niche position (range-shifting) and a wide niche breadth.

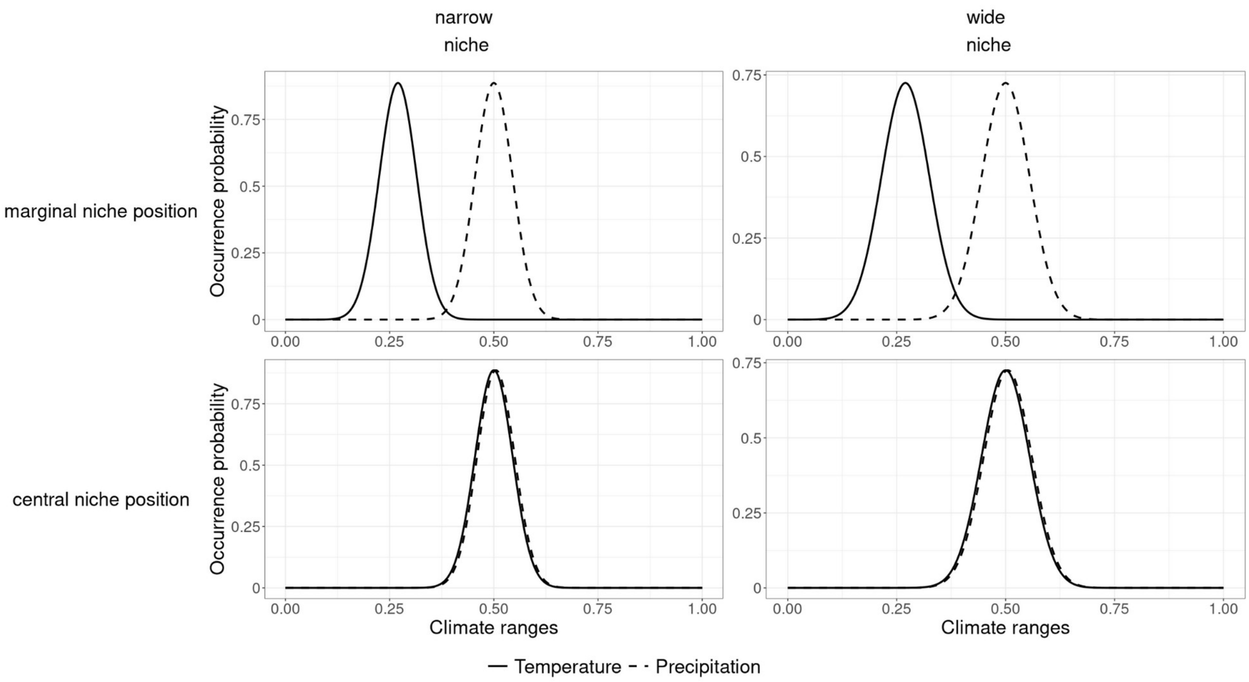


Figure S13: Response curves of the simulated niches of the virtual species for temperature and precipitation. Species with marginal, cold-adapted niche positions showed range-contracting under climate change. Species with central niche positions showed range-shifting under climate change.

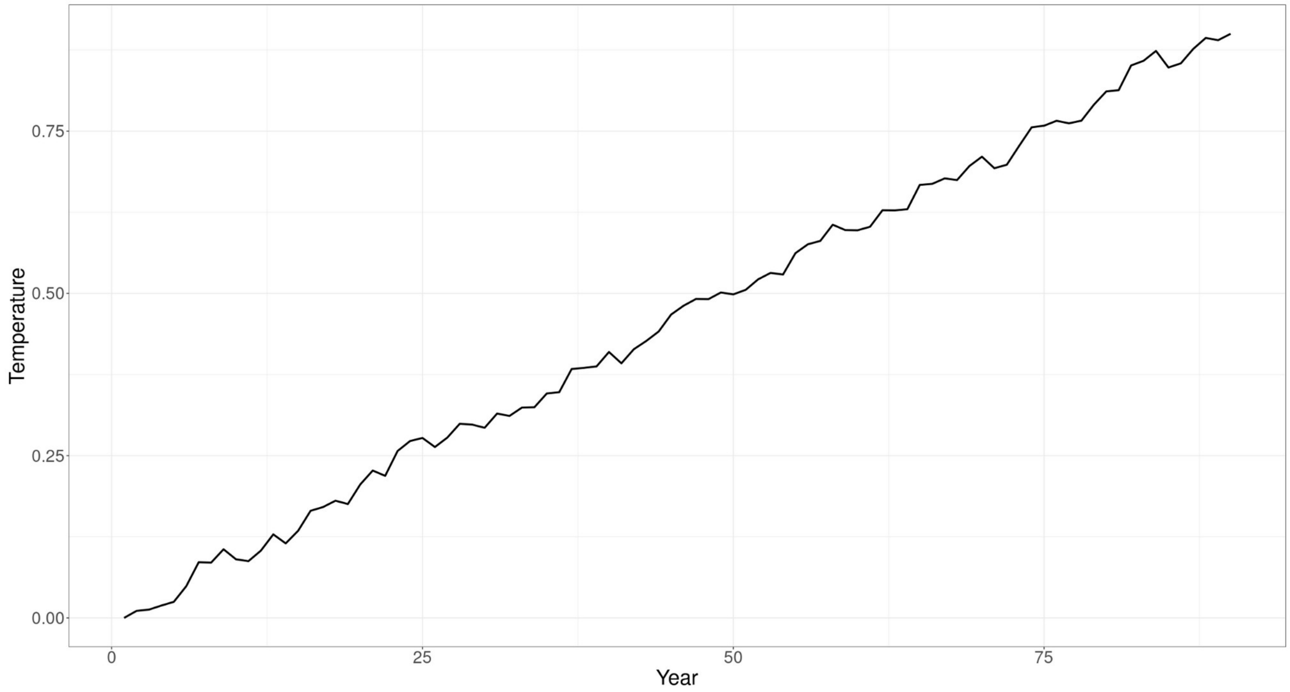


Figure S14: Modelled temperature increase over the 90 years of climate change.

Table S1: Results of the generalised mixed-model ANCOVA using an ordered beta regression with population size as the response variable and habitat loss, the four traits and the interaction between the traits and habitat loss as the predictor fitted to the ten selected replicated runs for all three landscapes and 16 virtual species. The landscapes were included as a random intercept. Depicted are the mean and the confidence values of the posterior distributions, the R-hat values, and Bulk and Tail Effective Sample Size (ESS) values.

| | | Mean | 95% CI | Rhat | ESS (Bulk, Tail) |
|---|---|---|---|---|---|
| Fixed effect | | | | | |
| | Intercept | 4.31 | 3.44, 5.48 | 1.00 | 1994, 1502 |
| | Habitat loss | -7.79 | -7.91, -7.67 | 1.00 | 3257, 4177 |
| | Niche position (range-shifting) | -3.22 | -3.31, -3.12 | 1.00 | 2891, 2454 |
| | Niche breadth (wide niche) | 0.57 | 0.52, 0.62 | 1.00 | 5343, 3605 |
| | Growth rate (fast) | 0.24 | 0.19, 0.29 | 1.00 | 5035, 5455 |
| | Dispersal (long) | 0.20 | 0.15, 0.25 | 1.00 | 4692, 3854 |
| | Habitat loss * Niche position (range-shifting) | 2.88 | 2.78, 2.98 | 1.00 | 3221, 4633 |
| | Habitat loss * Niche breadth (wide niche) | -0.32 | -0.39, -0.26 | 1.00 | 5414, 5577 |
| | Habitat loss * Growth rate (fast) | -0.11 | -0.18, -0.05 | 1.00 | 4792, 4937 |
| | Habitat loss * Dispersal (long) | 0.06 | -0.01, 0.12 | 1.00 | 4789, 3823 |
| Random effect | | | | | |
| | Landscape | 0.69 | 0.16, 2.46 | 1.00 | 1498, 896 |
